# Understanding the mechanism of monoADP ribosylation in OsSRT1 and its linkage to DNA repair system under stress conditions

**DOI:** 10.1101/2021.06.18.449075

**Authors:** Nilabhra Mitra, Sanghamitra Dey

## Abstract

The role of sirtuins in plants are slowly unraveling. There are only reports of H3K9Ac deacetylation by OsSRT1. Here our studies shade light on its dual enzyme capability with preference for mono ADP ribosylation over deacetylation. OsSRT1 can specifically transfer the single ADP ribose group on its substrates in an enzymatic manner. This mono ADPr effect is not well known in plants, more so for deacetylases. The products of this reaction (NAM and ADP ribose) have immense negative effect on this enzyme suggesting a tighter regulation. Resveratrol, a natural plant polyphenol proves to be a strong activator of this enzyme at 150 µM concentration. Under different abiotic stress conditions, we could link this ADP ribosylase activity to the DNA repair pathway by activating the enzyme PARP1. Metal stress in plants also influences these enzyme activities.

**Highlights:**

1. OsSRT1 can transfer a single moiety of ADP-ribose on itself as well as other nuclear proteins like histones H3 and H2A.
2. NAM, ADP-ribose and certain metal ions negatively regulate this ADP-ribose transfer.
3. ADPr of OsPARP1 and OsPARP2 links OsSRT1 to DNA damage repair pathways.
4. OsSRT1 positively regulates the activity of OsPARP1 by ADP ribosylating it.
5. On plant’s exposure to H_2_O_2_ (oxidative stress) and Arsenic toxicity, there is a link between the increased activity of the players of DNA damage repair system and overexpression of OsSRT1.

## Introduction

There are several studies showing the regulation and adaptation by plants in response to different stress conditions, both biotic and abiotic, in the environment they belong. This may be in the form of changes in histones as well as other non-histone protein modifications like acetylation, methylation, phosphorylation, and SUMOylation.

Sirtuins are class III HDAC superfamily members that require NAD^+^ for their catalytic abilities. Several alterations in the biological functions are the consequence of these modifications. Rice sirtuin, OsSRT1 belongs to class IV sirtuin family, having sequence identity with well-studied human SIRT6 (46 %) and SIRT7 (38 %). These human proteins are nuclear regulators against cellular stress conditions. OsSRT1 is instrumental in deacetylating H3K9Ac, which is linked with several metabolic processes in plants. One of the other post-translational modifications, ADP ribosylation is mostly linked to DNA repair, transcription and pathogenesis (1). This process can involve both mono as well as poly ADP ribose transfer from NAD^+^ molecule. In plants, this type of PTM is widely studied in PARP superfamily (2). There are certain members of the human PARP superfamily like PARP3, PARP7, PARP10 which are capable of mono-ADPr activity (3–5). But there is no data available for this process amongst plant PARPs. Mono ADPr is a reversible chemical reaction, used to regulate different cellular processes. The role of this activity in plants is not well understood except in its bacterial pathogenesis. Here the plant pathogens secrete effector molecules, which can ADP ribosylate the host proteins involved in plant immunity, in order to promote infection (6, 7).

In mammals it is mostly related to regulation under varied disease conditions (8). The ADP ribosylation of its substrates may have both positive and negative effect on them. SIRT1, SIRT2, and SIRT6 were shown to transfer mono-ADP-ribose moiety to BSA and histones *in vitro* (9, 10). SIRT4 has a negative effect on the mitochondrial metabolism by ADP ribosylating GDH (11). SIRT7, which is highly specific for the deacetylation of H3K18Ac, could ADP ribosylate itself as well as transcription factors like p53 and ELK4 (12). Both human SIRT6 and SIRT7 could influence the DNA repair process by ADP ribosylating PARP1 (12, 13). SIRT6 could also modify other nuclear proteins like repressor KAP1, resulting in repression of LINE1 transposable elements (14) and demethylase KDM2A to secure DNA repair (15). So, we can see that mono ADP ribosylation can affect the metabolic machinery at different cellular locations. This activity was also widely reported in sirtuins of several prokaryotes like *plasmodium falciparum* (16), *trypanosoma brucei* (17), *and salmonella* cobB (13). Yeast Sir2 also displayed this dual enzyme activity (19). There is a great possibility of crosstalk between different types of these modifications, say ADPr with acetylation (17) or ubiquitination with other PTM (20).

Mono ADP-ribosylation in plants by sirtuins have not been reported earlier. Among plant sirtuins, OsSRT1 has mostly been shown to be instrumental in deacetylating the H3K9Ac (21). Overexpression of this gene in rice plants resulted in enhanced tolerance to oxidative stress conditions, induced by paraquat. However, the molecular mechanism behind this action was not well understood. In this study, we could detect its dual enzyme action. Besides deacetylation, it can ADPr itself as well as other nuclear proteins. We could relate this activity to it providing tolerance to the rice plant under oxidative stress conditions. The regulation of this monoADPr activity in plants has also been discussed here.

## Material and Methods

### 2.1 Reagents

Restriction enzymes, Taq DNA polymerase and T4 DNA ligase were from New England Biolabs, USA. Primers were synthesized from IDT DNA, USA. All the chemicals of analytical grade were procured from Himedia, SRL chemicals and Sigma Aldrich Co., St. Louis, MO. Antibodies used in this study are anti His (sc-8036), anti PARP2 (PHY0218S), anti H3(ab18521), anti H4 (ab177840), anti PARP1 (ab110915), anti biotin (ab53468), anti-Poly ADPr (ab14459). rPARP1(ab123834) was purchased from Abcam, USA. Anti OsSRT1 against GST tagged core protein was custom synthesized from rabbit (Biobharati Pvt Ltd, India).

### 2.2 Homology Modelling of OsSRT1

Homology model of OsSRT1(483 aa) was generated in phyre2 server (22). The model was further minimized using modeller (23) to get most of the amino acid residues in the allowed region of Ramachandran plot parameters, as analysed in PROCHECK.

### 2.3 Cloning, overexpression and purification of OsSRT1 protein

The mRNA was extracted from the rice leaves using the kit (Himedia-HiPurA^™^ Plant and Fungal RNA Miniprep Purification Kit). This sample was converted into cDNA using Verso cDNA synthesis kit (Thermo Scientific, USA). The OsSRT1 gene was pcr amplified from the resultant cDNA using the gene specific primers. The *OsSRT1*gene was inserted in pET30a expression vector between *BamHI and EcoRI* restriction sites. The H134Y mutant was prepared using site directed mutagenesis protocol. All the DNA sequences of the OsSIRT1 constructs were checked by a DNA sequencing service.

The proteins were overexpressed in *E. coli* BL21 (DE3) strain. The cell culture was grown @ 37°C till its absorbance reached 0.7 and then induced with 1mM IPTG. This cell culture was further grown overnight @ 18°C. The cells were harvested by centrifugation (low temperature) at 4000 rpm for 30 min. The resultant pellet was resuspended in lysis buffer (50mM Tris pH 8.0, 300mM NaCl, 10% glycerol, 2mM DTT, 50mM imidazole). The cells were lysed using a sonicator (Hielscher). The cell debris were removed by a high speed (approx.14,000 rpm) centrifugation for 45 mins.

The filtered supernatant was loaded on a pre-equilibrated HisTrap FF column (GE Healthcare, USA). After thorough wash of the column, OsSRT1 protein was eluted in the buffer containing 200-250mM imidazole and further treated with DNAse (2μg/ml). The purified protein was run on SDS-PAGE and found to be more than 90% pure. The purified fractions were collected and desalted with 25mM Tris (pH 7.5), 150mM NaCl, and 2mM DTT for further experiments. Aliquots of desalted proteins were stored at −20 degrees. The final yield of OsSRT1 was approx. 0.8mg per gram of cell.

IR64 and Swarna variety of rice seeds were first surface sterilized and laid on cotton sheets for germination in dark for 48 hours. Then it is transferred to plant culture bottles with MS solution and left at 25°C with 10h light and 14h dark cycles for about 3-4 weeks till 3 leaf stage. For different stress treatments, three-week-old rice seedlings growing in MS medium were transferred to MS medium supplemented with 500 mM NaCl or 50mM AlCl_3_ for 2 days or plants were sprayed with 10% H_2_O_2_ for the same duration. For dehydration stress conditions, the 3 week old plants were taken out of MS media for 2 days. All the plant samples were flash frozen and stored at − 80°C.

### 2.4 Mono ADP ribosyl transferase activity of OsSRT1

The ADP ribosylation reaction was performed at 25°C for 1 hrs. Buffer composition for the reaction was 50 mM Tris pH 7.5, 150 mM NaCl, 10 mM DTT and 75 μM NAD^+^ and 40 μM 6-biotin-17-NAD (Bio-NAD^+^, R&D, Minneapolis, USA). The reaction mix containing 0.8μg purified OsSRT1 were then resolved using 12% SDS-PAGE and the immunoblot analysis was done by initially blocking the nitrocellulose membrane in TBST for 30 mins. Rabbit monoclonal anti-biotin antibody (ab19221) was used to detect the biotinylated ADP ribose moiety of the protein. This was followed by incubation with HRP conjugated secondary antibody and developed with ECL solution (BioRad). This experiment was further repeated with recombinant Histones (NEB) (0.5µg) and PARP1 (5μg) as substrates for OsSRT1. ADPr transferase activity has also been examined in presence of 0.5 µM concentration of various salts (CuSO_4_, ZnCl_2_, FeCl_3_, MgCl_2_, CaCl_2_ and MnCl_2_). All the experiments were carried out in triplicates. ImageJ software (NCBI) was used for the analysis of the band intensity in the blots (24). For dot blot analysis, 6nM OsSIRT1 was mixed with varied concentrations of Bio-NAD^+^ (0-100 μM). For the determination of kinetic parameters of H3 ADPr reaction, constant conc. of H3 (3.3nM) was added to the same kind of reaction mix

### Dot Blot Analysis of OsSRT1 enzyme activity

The reaction mixtures were prepared and carried out as mentioned above. The finished reactions were desalted to remove the unused Bio-NAD^+^ or other buffer constituents. The reaction mix was then dot blotted on a nitrocellulose membrane and air dried. The blots were developed as in western blot protocol. The standard curve of Bio-NAD^+^ was also prepared for the said reaction. The intensity of the dots were analyzed using ImageJ software (NCBI) (24) and Microsoft Excel. Saturation kinetics using Non-Linear regression (Michaelis Menten Model) was prepared using GraphPad Prism version 9.1.1 (San Diego, USA).

## Result and Discussion

### (1) Structural features of OsSRT1

Using the available coordinates of human homologs and *de novo* modeling, we have prepared the model of OsSRT1 using phyre2 server (22). The model clearly delineates the two areas: one for binding the NAD^+^ (Rossmann Fold) and the other is for the Zn^2+^. In addition, there is a huge C-terminal region (290-483) not present in the other classes of plant sirtuins, forming a separate domain. The function of this domain is not quite clear, but may be speculated as a regulatory site or accessory protein binding site for this enzyme. The peptide/substrate binding loop (179-188) contains the motif WEDAL unique to this class IV. This motif is present in between the Rossmann Fold and Zn binding domain. The two loop regions harbor the four Cys (Cys^142^, Cys^145^, Cys^167^ and Cys^172^) residues for tetrahedral coordination with Zn^2+^ ion. The active site of OsSRT1 is further lined by several loops containing the conserved motifs like GASISTSSGIPDFR, QNVD. All these structural features contribute towards its substrate specificity. **(Fig1)**

**Fig1.**
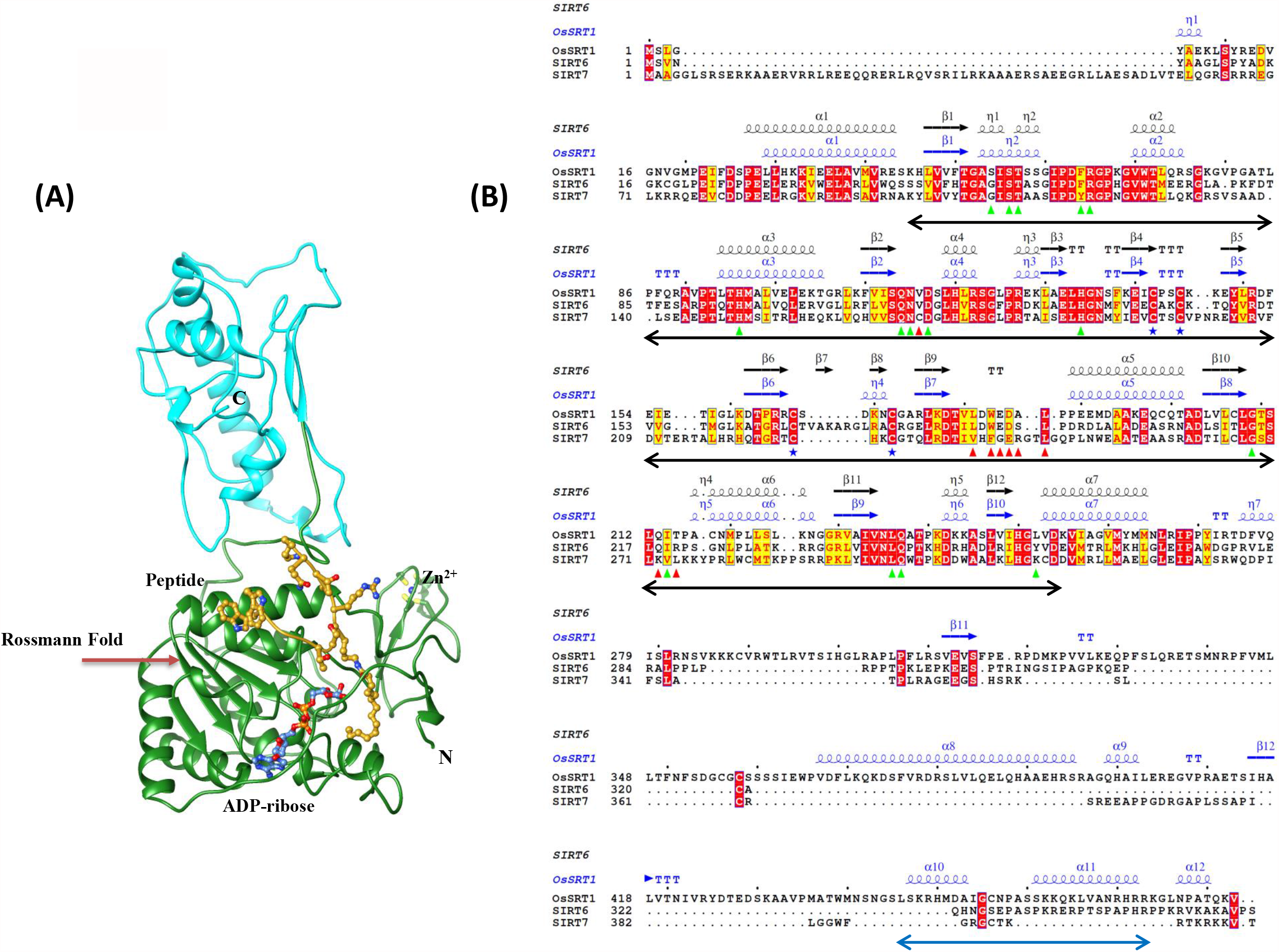
Structural analysis of OsSRT1 enzyme: **(A) Homology model of OsSRT1 in complex with the substrates:** Figure shows the cartoon depiction of the OsSRT1 model in complex with the ligands ADP-ribose and H3 peptide in ball and stick form. It clearly shows the two adjacent regions where the two ligands bind at the active site. Away from the active site, Zn^2+^ (blue) is present in coordination with four Cys residues (yellow). The extra C-terminal region is shown in cyan color. (B) **Sequence analysis of class IV sirtuin members:** Sequence alignment of OsSRT1(B8ARK7) with its human homolog SIRT6 (Q8N6T7) and SIRT7(Q9NRC8). The residues which line the NAD^+^ binding region are shown in green triangles, peptide binding residues with red triangles, cysteines involved in zinc binding are shown with blue stars. The catalytic base His 134 is shown with a red circle. The underlying black arrow demarcates the catalytic core region. The secondary structure depiction is based on the homology model coordinates. The underlined blue arrow at the C-terminus depicts the NLS based on cNLS mapper analysis. The figure is prepared using ESPript 3.0 (1).

### (2) Localisation and tissue specific expression of SRT1 protein in *oryza sativa indica*

Western blot analysis was performed using the custom made anti-OsSRT1 antibody to detect its presence in different organelles of *Oryza sativa indica* sample. We have extracted the organelles (nucleus, cytoplasm and mitochondria) from the 2-week-old rice leaf cuttings. Our studies have detected this protein solely in the nucleus under normal plant conditions. This data is supported by its localization in nucleus using GFP tagged SRT1 in onion epidermal tissues (25) and protein sequence analysis in MultiLoc2 server. Further tissue analysis showed the expression of SRT1 protein in all the major tissues like stem, leaves and roots. **(Fig2a and b)**

**Fig2.**
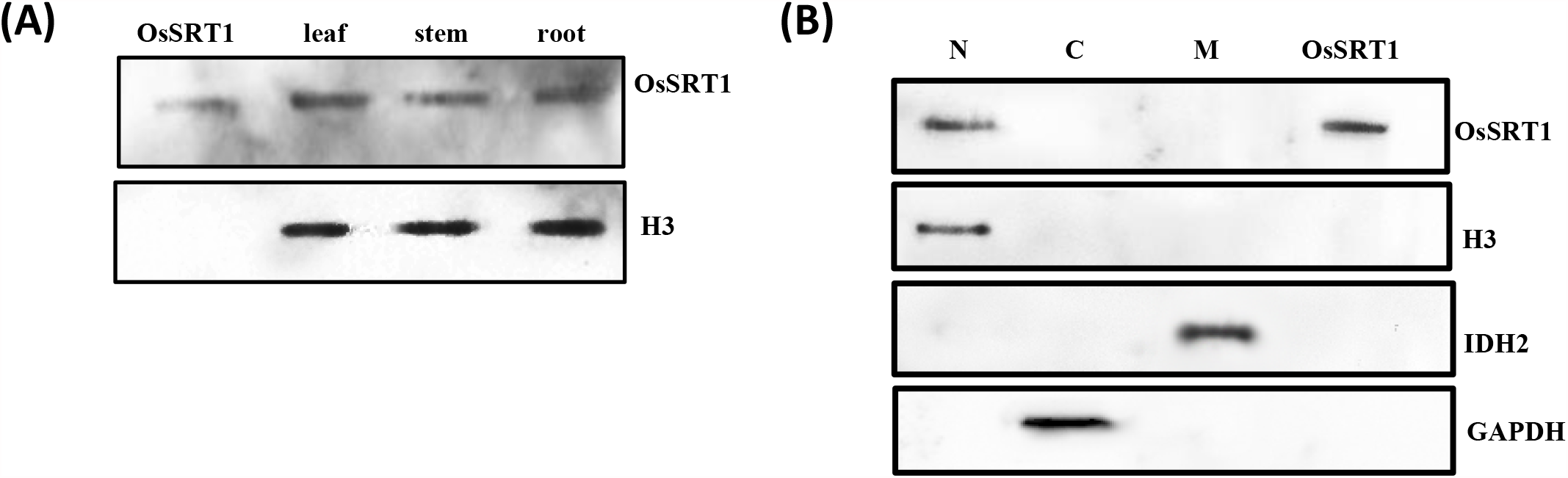
Protein Expression of OsSRT1 in *Oryza sativa indica*: (A) Western blot analysis of the major tissues (Leaves, stem and roots) of two week old rice plant to detect the presence of OsSRT1. (B) Cellular localization of this protein suggests its presence in the nucleus. Here organelle specific markers are used to show the purity of the fractions (nucleus N, cytoplasm C and Mitochondria M). For both these experiments, immunoblots were developed with the anti OsSRT1 antibody to detect its presence.

### (3) OsSRT1 shows NAD^+^ dependent auto ADPr activity

As there is evidence of H3K9Ac deacetylation by OsSRT1(21), we were interested in dissecting its other enzymatic abilities. In our studies, we found that the recombinant protein (1µg) can fully ADP ribosylate itself within 1hour time (**Fig3a)**. In plants auto ADPr activity of deacetylases has not been reported yet. In our experiments we find that OsSRT1 is capable of transferring the ADPr moiety on itself (**Fig3b)**. For this it requires NAD^+^ in its reaction. The H134Y mutant is not able to ADP ribosylate itself suggesting the requirement of this residue for this catalysis. We further wanted to know whether OsSRT1 transfers a single or multiple units of ADPr on itself. We find that this enzyme is only capable of transferring the single moiety of ADPr. In this respect, we have used recombinant PARP1, which is capable of transferring multiple copies of ADPr moieties, as a positive control for this study **(Fig 3c)**. Once we have established that OsSRT1 is a mono ADPr, we looked for the amino acid residues which can get modified. Chemical stability assay of this enzyme’s auto ADPr revealed that the transfer is mostly targeted to Arg residues of the substrate. In this experiment, reagents like HgCl2, NaCl and NH2OH were used to detect the cleavage of covalent linkage of ADP ribose moiety with Cys, Lys and Arg residue, respectively (26–28). The blot showed a faint biotin band for the reaction mixed with NH2OH in comparison to the other reagents. This indicates the hydrolysis of the respective Arg linkage with ADPr **(Fig 3d)**. Interestingly, its human counterparts, SIRT6 is specific for Lys residues (13) and SIRT7 is specific for Arg and Cys (12).

**Fig3.**
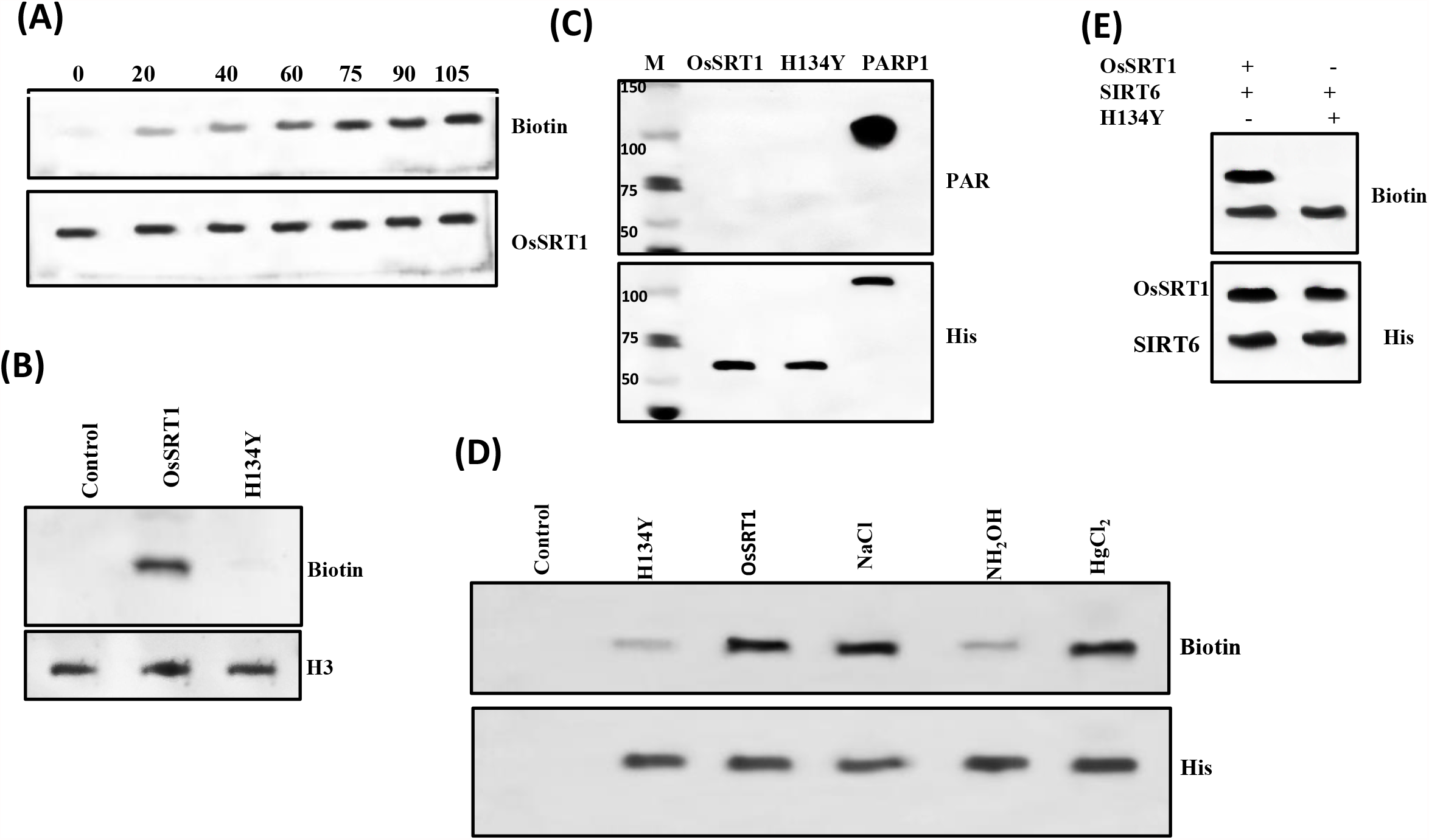
OsSRT1 shows monoADP ribosylation activity: Here we have used H134Y mutant and empty vector (control) as the negative control. **(A)** Timeline of monoADPr reaction of OsSRT1. OsSRT1 can be auto ADP ribosylated in a time dependent manner using leaf nuclear extract. **(B)** The blot showed that the OsSRT1 can ADP-ribosylate itself using substrate Bio-NAD^+^ (40µM). Its catalytic (H134Y) mutant did not show this transfer as evident from no biotin band in this lane. These reaction mix containing nuclear extract were carried out at 25°C for 1 hr. Histone H3 was used as loading control in lower panel. **(C)** The anti poly ADPr (PAR) western blot demonstrated that OsSRT1 is not capable of transferring multiple units of ADP ribose on itself. Here, we have used the H134Y mutant and PARP1 as negative and positive control, respectively. **(D)** Chemical stability analysis of auto-ADP ribosylation of OsSRT1 suggests Arg as the site of ADP-ribosylation. Purified OsSRT1 was incubated with Bio-NAD^+^ in the reaction buffer at 25°C for 1 hour. After that, the reaction mix was divided in equal aliquots and different hydrolyzing agents (0.5mM) were added to it and kept for another 2 hour at the same temperature. The reaction was stopped by adding sample dye. The reaction mixture was then resolved on SDS PAGE gel and anti biotin immunoblot was carried to detect the presence of biotin. **(E)** OsSRT1 ADP ribosylates in an enzymatic manner: Here OsSRT1 and H134Y mutant were used as substrates for ADPr transfer reaction in presence of SIRT6. Human SIRT6 has the capability to transfer ADPr residue. This western blot was performed to check for the non-specific binding of ADPr with the rice protein. This enzyme which also has the ability to self ADP ribosylate, we see a biotin band in the lanes. The H134Y mutant, which cannot hydrolyse NAD^+^, did not show a non specific binding of ADPr moiety with the mutant (Lane2). Thus, SIRT6 is unable to transfer the ADPr moiety on the mutant.

### Enzyme kinetics

As sirtuins are mostly considered as NAD^+^ dependent deacetylases so there have been always doubts regarding the existence of this ADPr activity. It is mostly thought to be a result of a side reaction of deacetylation. After the hydrolysis of NAD^+^ by sirtuins, there is a possibility that the excess ADPr moiety present in the medium can get nonspecifically bound with the amino acid residues of a protein. To check this fact, we performed an experiment using SIRT6 as a control. This enzyme is also capable of mono ADPr activity and can hydrolyse the NAD^+^ into NAM and ADPr (10, 13). Here, we did not find the transfer of ADPr moiety on the OsSRT1 H134Ymutant, which itself is not capable of hydrolysing the NAD^+^ molecule (**Fig3e)**. So, we think that the observed mono ADPr by OsSRT1 follows an enzymatic reaction mechanism and the transfer is specific. The Kcat and Km values of the auto ADPr reaction of OsSRT1 is 72±3 min^-1^ and 37.44±4 µM, respectively. Thus, it is clear that OsSRT1 favors the ADPr reaction more than deacetylation (Kcat= 4.5±2 min^-1^) as the rate of ADPr is 20 times more than deacetylation. **(Fig4c)**

**Fig4.**
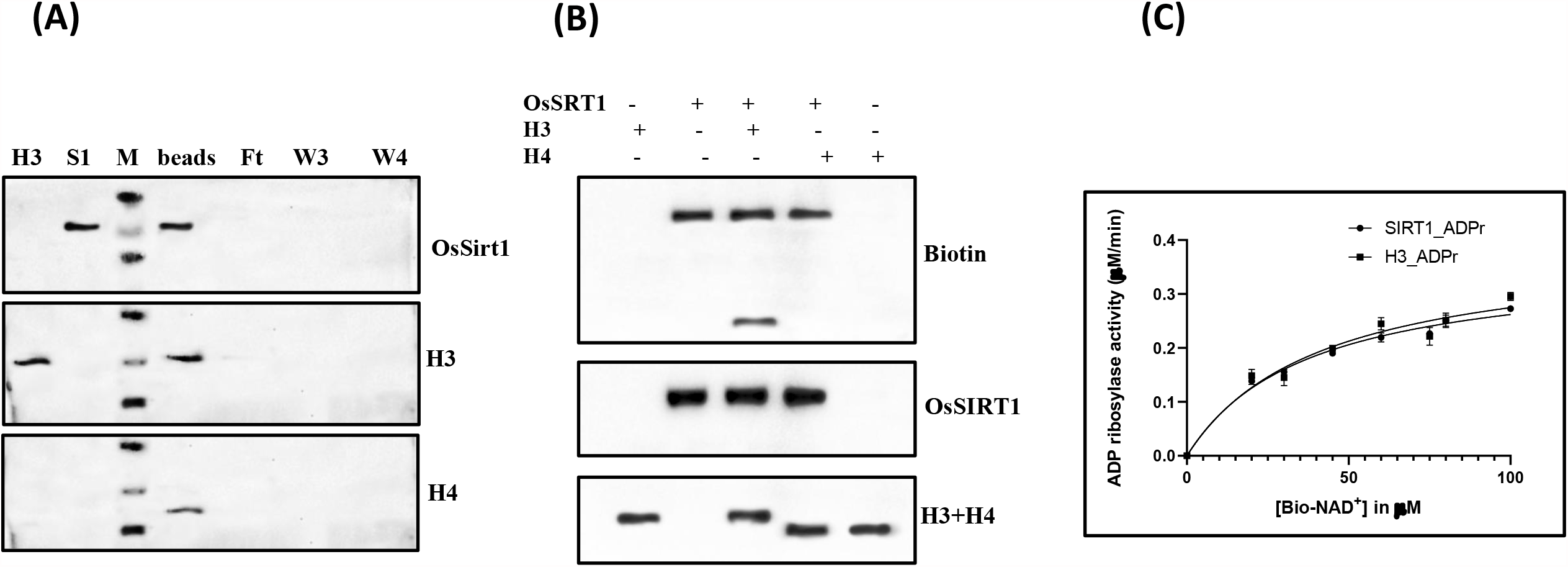
Histones as target of OsSRT1 MonoADP ribosylation: **(A) Interaction of OsSRT1 with histones H3 and H4**. Ni pull down assay was performed to detect the interaction of His-tagged OsSRT1 with histones extracted from the nuclear fractions of rice leaves. The resultant protein complexes pulled down by Ni-NTA beads were resolved on SDS-PAGE and presence of H3 and H4 in these complexes was detected by histone specific western blot. Initial flowthrough (Ft) and final washes (W3 and W4) of this assay were also run on the gel. Here, H3 and H4 themselves did not nonspecifically bind to the Ni beads. **(B) MonoADPr analysis of OsSRT1 with H3 and H4**. Western blot analysis showing the ADPr transfer reaction of OsSRT1 with histones H3 and H4 as substrates along with Bio-NAD^+^. Reactions were carried out with and without recombinant histones. OsSRT1 can hydrolyse the NAD^+^. The resultant free ADP ribose does not bind non specifically with the histones as evident from the lane containing H4 (lane 4). The blot shows biotin band for OsSRT1 (self ADP ribosylation) and histone H3. This experiment shows that OsSRT1 can ADPr histone H3. The loading control, using anti H3+H4 western is shown in the lower panel. **(C) Saturation kinetics for the OsSRT1 monoADPr activity**. Auto ADPr transfer reaction of the OsSRT1(6nM) using varied concentration of Bio-NAD^+^ (0-100 µM). The same reaction in presence of histone H3 (3.3nM) was also done. The plots were calculated using Graphpad Prism 9.1.2 (MM Model). The error bar depicts the S.D.; n=3. Further kinetic parameters (K_m_ and K_cat_ values) were calculated using these plots to understand the dual activity of OsSRT1.

Besides its self-modification by ADPr, we also looked for other nuclear targets which can get ADP ribosylated by OsSRT1. Ni pull down assays with the leaf extract showed that this enzyme can directly interact with both histones, H3 and H4 but only modifies H3 by ADP ribosylation. **(Fig4a&b)**. Among other histones, OsSRT1 can also transfer a single copy of ADP ribose on H2A as a target **(FigS1)**. In these experiments, both the histones (H3 and H2A) were not modified by other enzymes in the leaf extract as evident from the lanes containing the H134Y mutant and empty vector as a control. We have also shown that the ADPr transfer on H3 is not non-specific **(FigS2a)**. All these data suggest that this histone modification is brought about by OsSRT1. In comparison histones are not the substrates for its human homolog Sirt7 mono ADPr activity. SIRT6 can ADP ribosylate H3 as a target **(FigS2b)**.

In this study we show for the first time that a plant sirtuin OsSRT1 is a real mono-ADP-ribosyltransferase that catalyses both auto- and hetero-modification by transfer of a single ADP-ribose moiety from NAD^+^. It prefers this activity more than H3K9 deacetylation based on its kinetic parameters.

### (4) Modulation of OsSRT1 ADP ribosylation activity

Once we detected this activity in OsSRT1, we wanted to understand its regulatory mechanism in cells. Sirtuins depend on NAD^+^ for their enzyme activity but also need Zn^2+^ ions for structural stability. In absence of Zn^2+^ ion, the overall structure of sirtuin may collapse (29) or excess Zn^2+^ ion can also inhibit its catalysis (30). The role of metal ions in mono ADPr reaction is not well studied, though high concentration of these metals can have toxic effect on plants (31). In our experiments, the endogenous Zn^2+^ ion was sufficient for the ADPr reaction of OsSRT1. We have tested several metal ions to detect their effect in this reaction. 0.5 µM concentrations of Cu^2+^, Mg^2+^, Mn^2+^ and Ca^2+^ had great inhibitory effect on OsSRT1 ADPr activity. Same concentration of Fe^3+^ ions also hampered this effect to some extent (approx. 30%). **(Fig 5a)**. The molecular mechanism for this metal inhibition on OsSRT1 activity in plants is not clear. There is a possibility of presence of a regulatory site in OsSRT1 where these metal ions can bind and bring about this modulation (allosteric effect). Excess metal ions can also unfold the proteins as they mostly damage the targeted proteins in plants during toxicity.

**Fig5.**
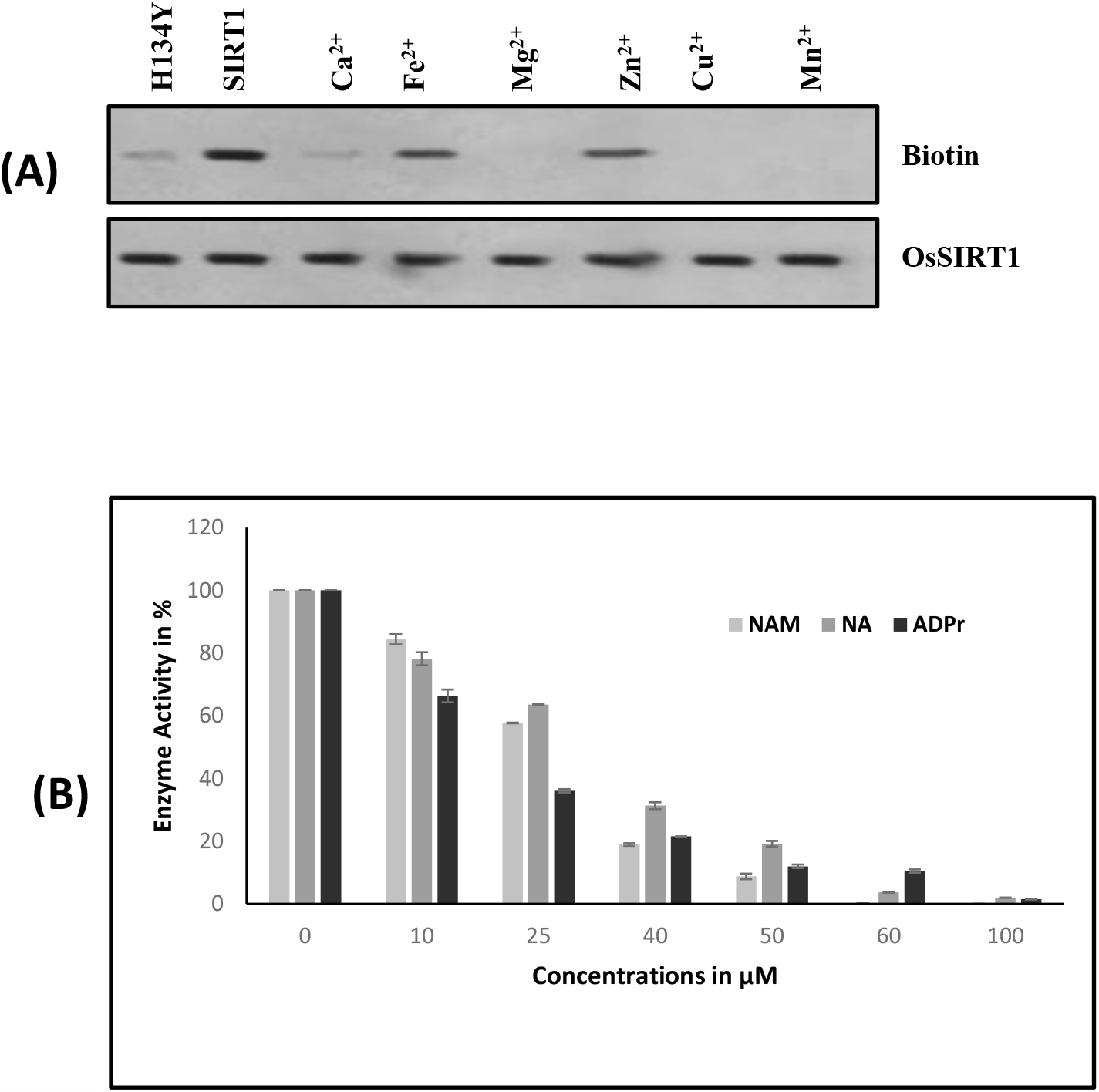
Modulation of ADPr transfer activity of OsSRT1: For western blot analysis, anti-biotin antibody was used. **(A) Effects of different metal ions on ADPr transfer activity of OsSRT1**: This figure shows the effect of different cations on the auto ADPr activity of OsSRT1. several metal ions like Ca^2+^, Fe^3+^, Mg^2+^, Cu^2+^ and Mn^2+^ (0.5mM) were added to the reaction mix and checked for their effect in western blots. H134Y mutant was used as a control. **(B) Product inhibition of OsSRT1 activity:** A bar diagram showing the comparative analysis of NAM, NA and ADP-ribose effect on auto-ADPr transfer of OsSRT1. Western blot analysis was performed to measure the effect of product nicotinamide (NAM) on the auto-ADPr modification of OsSIRT1. The reactions were performed in triplicates and the error bar shows the S.D. Auto-ADPr transfer activity was detected by anti-biotin antibody. The reactions were carried out in presence of increasing concentration of NAM, NA and ADP-ribose (0-100µM).

Product inhibition is another way of regulating any enzyme function. In the OsSRT1 reaction, along with deacetylated lysine, nicotinamide (NAM) and 2’O-acetyl ADP-ribose (OAADPr) are the products that are released on hydrolysis of NAD^+^. We have performed the enzyme reaction in presence of increasing concentration of NAM (0-100μM) to study its effect on ADP ribosylation. IC_50_ value of NAM inhibition for ADPr reaction was 26±2 μM. Similarly, nicotinic acid (niacin, NA), lacking the amide group of NAM, had IC50 value of 32±4 μM for OsSRT1 ADPr reaction. Since the cellular NAM concentration is estimated to be between 10-150 μM (32) so OsSRT1 can be regulated by physiologically relevant NAM concentration. Interestingly, the auto modification of OsSIRT1 is also quite sensitive to the other product analog ADP-ribose with IC_50_ value of 18±4 μM. Thus, product inhibition seems to be an effective way of sirtuin’s regulation in plants. Also, this enzyme is quite sensitive to this type of regulation in comparison to its human counterparts (**Fig 5b)**.

In addition, Resveratrol, a plant polyphenol and also well-known sirtuin activator showed positive effect on the ADPr activity of OsSRT1 at the concentration of 150µM **(FigS3**). SIRT6 and SIRT7 activities were not affected by it (12). Resveratrol is a natural polyphenol mostly produced in grapes, peanut and few other plants by stilbene synthase of phenylalanine/polymalonate pathway. So, if transgenic plants with this pathway is created in rice plants that can upregulate the SIRT1 activity, making the plants more tolerant against stress conditions. This effort will be quite beneficial to the crops and increase their yield. Earlier, these types of transgenic rice plants have been created for the betterment of human disease conditions (33).

Under different stress conditions in plants, the auto modification of OsSRT1 may also affect the interaction of this enzyme with its regulatory partners. Thus, modulating the role of this enzyme.

### (5) Biological role of OsSRT1 ADPr activity in plants

The physiological significance of this enzyme in plants is still unfolding. Here, we were further interested in corelating this modification with its cellular functions. In 2007, Huang *et al*. analyzed the effect of RNA interference (RNAi) of *osSRT1*gene using the ChIP assay. Downregulation of this gene increased the acetylation of histone H3K9 and reduced its demethylation on transposable elements and hypersensitive response (HR)-related gene promoters. This caused an increased H_2_O_2_ production, resulting in DNA damage and cell death in plants. Overexpression of *osSRT1* gene increased the plant’s tolerance to this oxidative stress. However, the mechanism behind this OsSRT1 action and its link to molecular machinery was not clear (21).

To answer this question, we performed Ni pulldown experiment of His tagged OsSRT1 with the nuclear extract of rice leaves. Interestingly, we found that among several other proteins, OsSRT1 interacted with the players involved in DNA damage and repair pathways i.e. OsPARP1 and OsPARP2 (34–36) **(Fig6a)**. OsSRT1 was also capable of transferring the biotinylated ADPr to these proteins **(Fig6b)**. In turn, OsPARP1 cannot poly ADPr this rice sirtuins (data not shown). This finding is quite relevant as PARP1 activity also increases (approx. 75%) on ADP ribosylation by OsSRT1**(FigS4)**. To explore the role of ADPr in plants under stress, they were exposed to various abiotic stress conditions like salinity, dehydration and high concentrations of chemicals like AlCl_3_, As, H_2_O_2_. Under the conditions of excess H_2_O_2_, Arsenic toxicity and dehydration, the expression levels of OsSRT1, PARP1, and PARP2 got upregulated **(Fig6c)**. We can see a correlation between the increased expression of OsSRT1 with the increase in the poly ADPr activity of PARP1**(FigS5)**. Here, we can speculate that under oxidative stress conditions, OsSRT1 can invoke the DNA repair system when there is DNA damage due to toxicity or oxidative stress. There might be some relationship between the increased expression of OsSRT1 with the DNA repair pathway.

**Fig6.**
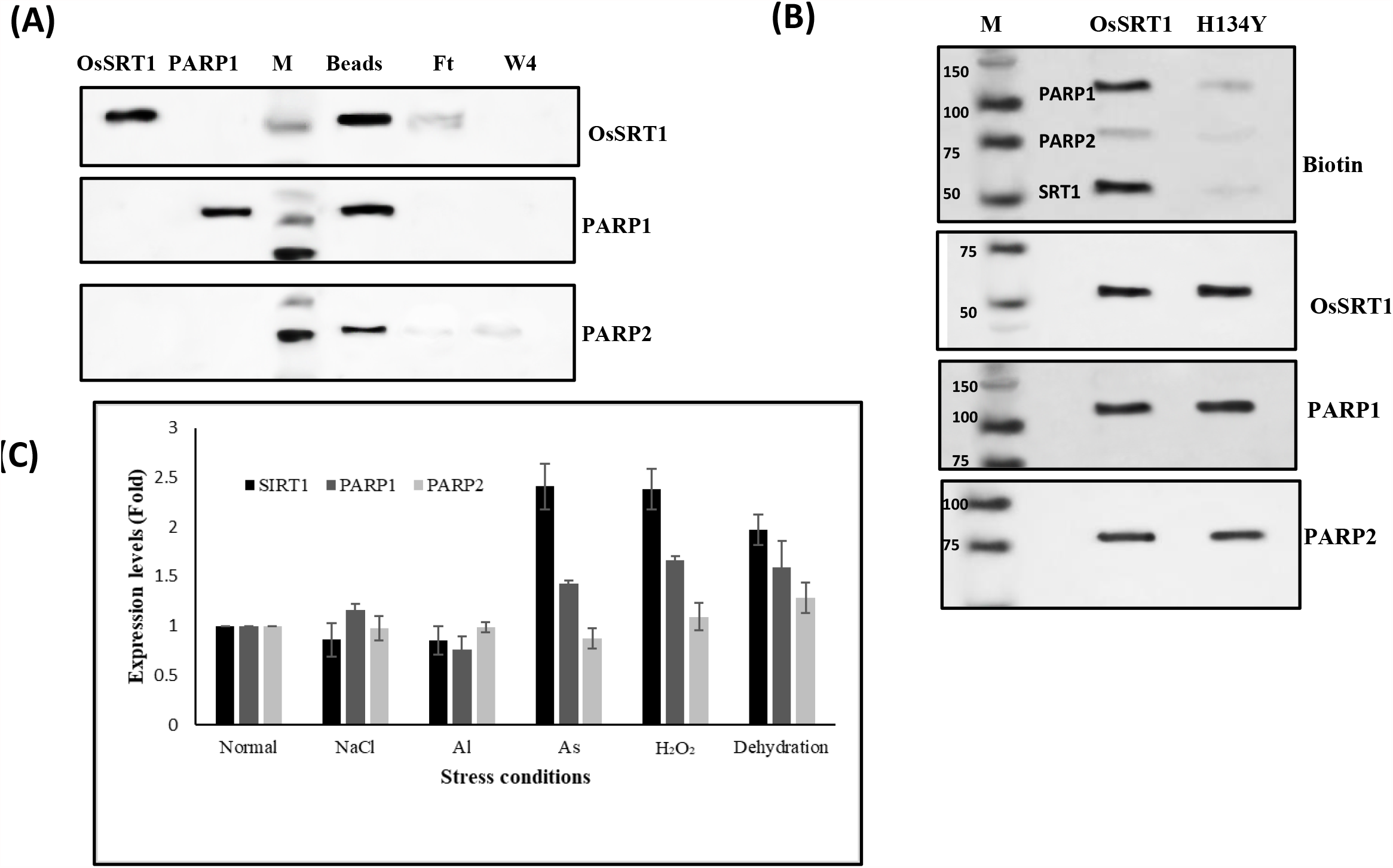
Biological roles of mono ADPr transfer activity of OsSRT1: **(A) Detection of nuclear substrates from rice leaves**. Ni pull down experiment of OsSRT1 with nuclear extract of rice leaves showed the interactions with OsPARP1 and OsPARP2 as detected by protein specific antibodies. **(B) Plant PARPs are targets of ADP ribosylation by OsSRT1:** Recombinant OsSRT1 is able to transfer the biotinylated ADP-ribose on OsPARP1 and OsPARP2. The nuclear extract was incubated with recombinant OsSRT1 in an ADP ribosylation reaction mix containing Bio-NAD^+^ for 1 hr at 25°C. The reaction was stopped by adding sample dye, boiled and resolved on SDS-PAGE. It was then transferred on nitrocellulose membrane. The blot was probed with anti-biotin antibody to detect the presence of ADP ribose on respective proteins. Here the negative control is the empty vector or H134Y mutant. The loading control is shown in the bottom panel with blots developed with protein specific antibodies. **(C) Expression analysis of OsSRT1, OsPARP1 and OsPARP2 proteins under different stress conditions**. The nuclear protein extract was prepared from the leaves of the plants exposed to the different stress conditions as mentioned in the figure. Western blot analysis was performed using the specific antibodies (OsSRT1, PARP1 and PARP2). Histone H4 in the nuclear extract was used as the internal control. All the data were normalized before plotting. The expression of OsSRT1 and OsPARP1 increased under excess As, H_2_O_2_, and dehydration. OsPARP2 showed shuttle differences in expression for these stress conditions.

The expression level of PARP1 and PARP2 in arabidopsis have also been found to be induced on DNA damage under oxidative stress condition (34). With elevated OsSRT1 expression, ADPr activity of PARP1 will also increase and this will activate the PARP1 activity further under stress conditions to stop the DNA damage. Similarly, histone ADP ribosylation in response to DNA damage is quite prevalent in nature. In addition, histone ADPr is also linked to DNA repair pathways. Though much is not known about this in plants.

## Conclusions

OsSRT1 is almost ubiquitously present in the major tissues of the plant. In this study, we are reporting the mono ADPr activity of OsSRT1 for itself as well as other nuclear proteins in plant. This sirtuin possesses dual enzyme activity similar to its human homologs. We also see a similar trend in case of OsSRT1 with a weaker deacetylase activity compared to its ADPr activity. The structural features of this plant sirtuin is highlighted with a separate C-terminal domain, whose function needs to be explored. The ADPr activity is also very sensitive to the products of the enzyme reaction, NAM and ADPr. NAM can play a major role in balancing the excessive breakdown of NAD^+^ in response to stress conditions, thus maintaining the metabolite homeostasis in cells. Metal ions like Cu^2+^, Mg^2+^ and Ca^2+^ have an inhibitory effect on this action suggesting a big impact on epigenetic regulation in plants under metal stress. Resveratrol, a plant polyphenol has a positive effect on this enzyme action. Thus, there is possibility of production of transgenic plants overexpressing this molecule, which can make these plants more tolerant to the abiotic stress conditions.

ADP ribosylation is known to play a role in regulation of DNA damage and repair pathway (1). This kind of ADPr reaction may be reversible process as there is existence of eraser proteins in plants (37). While searching for its biological relationship in plants, we found that OsSRT1 can interact directly with histones and the components of DNA repair pathway as evidenced from the Ni pull down experiments. Both PARP1 and PARP2 proteins are involved in DNA repair process in plants (34, 38). This mono ADPr action is involved in regulating OsPARP1’s activity in a positive way. Furthermore, under metal stress conditions, when the OsSRT1 activity is inhibited, we also see a decrease in activity of these enzymes. This reduction in its PAR activity may be a direct effect of the metal ions or may be due to the OsSRT1 action. But this suggests that metal toxicity in soil can easily hamper the DNA repair mechanisms in plants.

In addition, it is known in mammals that histone ADP ribosylation is also induced by DNA strand breaks (39, 40). These effects play an important role in metabolism and disease conditions. Similarly, we may speculate here that OsSRT1 action on histones H3 and H2A may be in response to the DNA damage in plants under stress conditions.

Overall, this ADPr study reveals an important aspect of OsSRT1 action in chromatin regulation in plant stress responses. With the elucidation of mono ADP ribosylation in plant proteins, it opens up the avenues to understand the mechanism of adaptation or tolerance under unfavorable conditions.

There are still several questions which needs to be further enquired: How is the dual activity regulated in sirtuins harboring both enzymatic activities? What makes one activity more important than the other for a specific substrate? Under what circumstances in plants, OsSRT1 ADP ribosylates itself or is it a constitutive process. It needs to be seen what are the other proteins which are recruited during DNA damage in plants because of ADP ribosylation. As epigenetic regulations also involve covalent modifications in DNA or RNA, does OsSRT1 has the capability to ADP ribosylate these molecules? With this ability in its arsenal can plant also withstand the biotic stress created by bacterial or fungal pathogens?

## Abbreviation

ADPr: ADP ribose/ADP ribosylation
HDAC: Histone deacetylase
PTM: Post translational modification
PARP1: Poly ADP-ribose polymerase 1
PARP2: Poly ADP-ribose polymerase 2
GDH: glutamate dehydrogenase
NAM: Nicotinamide
NA: Nicotinic acid

## Supporting information

**Figures S1-S5**

## Acknowledgement

This work was funded by Department of Science and Technology, Govt Of India (CRG/2019/003037) and FRPDF scheme, Presidency University, Kolkata, India.

## Supplementary figures

**FigS1.**
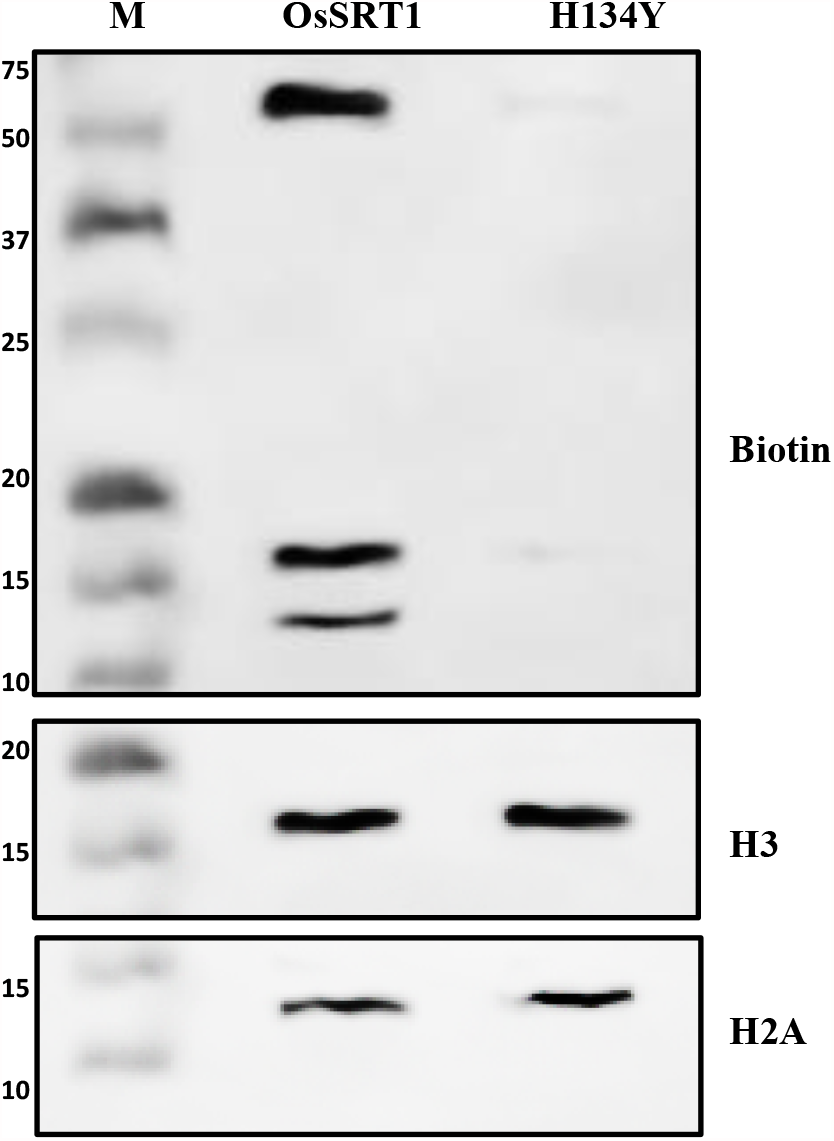
OsSRT1 can monoADP ribosylate histones: Anti biotin western blot analysis showing the result of OsSRT1 ADPr reaction with leaf extract. H134Y mutant was used as a negative control. The lower panel shows the loading control in the form of H3 and H2A bands developed with the protein specific antibody.

**FigS2.**
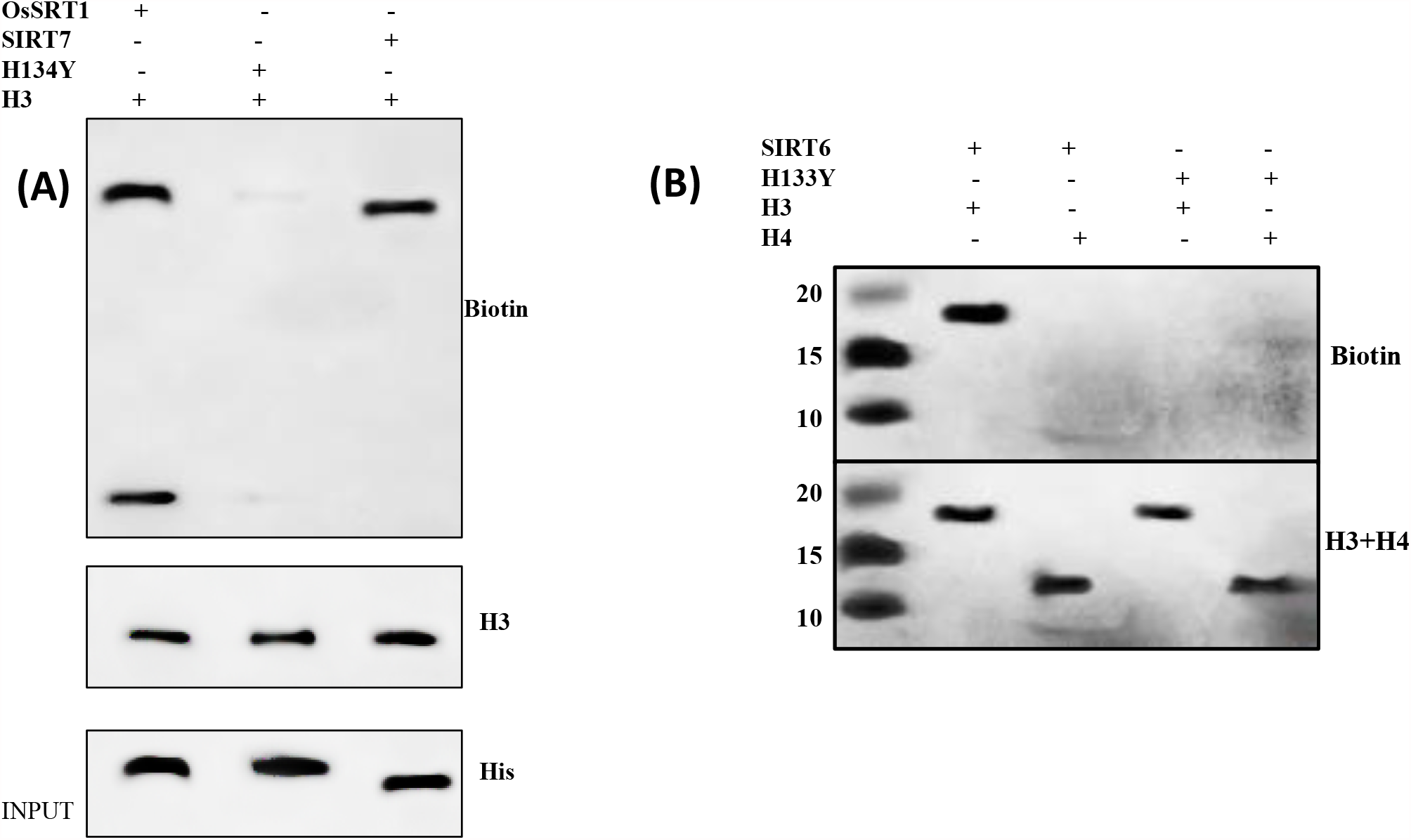
**(A)** Histone H3 modification in presence of OsSRT1 and its homolog SIRT7. OsSRT1 mutant did not ADP ribosylate H3. **(B)** SIRT6 can ADPr histone H3 but not H4. The lower panel shows the loading control developed with H3+H4 antibody.

**FigS3.**
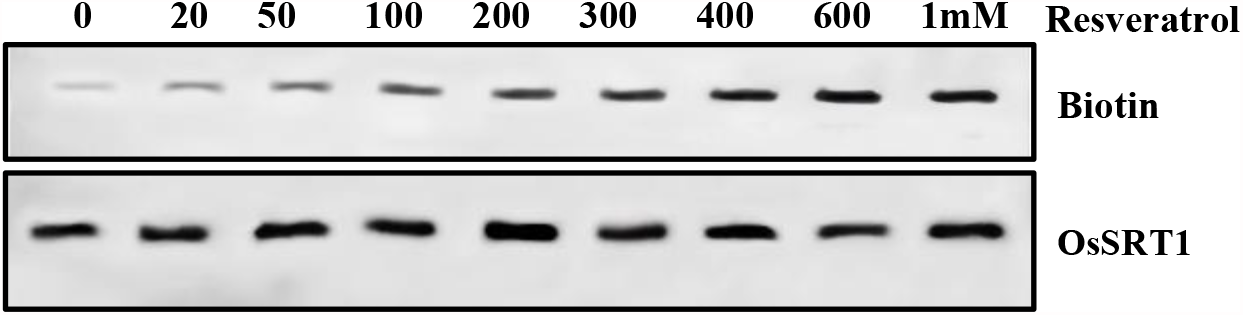
Effect of polyphenol Resveratrol on the OsSRT1 monoADPr activity: Anti biotin western blot shows the effect of resveratrol on the auto modification of OsSRT1. The activator was added in the reaction mix in the range of 0-1000 µM. The lower panel shows the loading control blot developed with anti-OsSRT1 antibody.

**FigS4.**
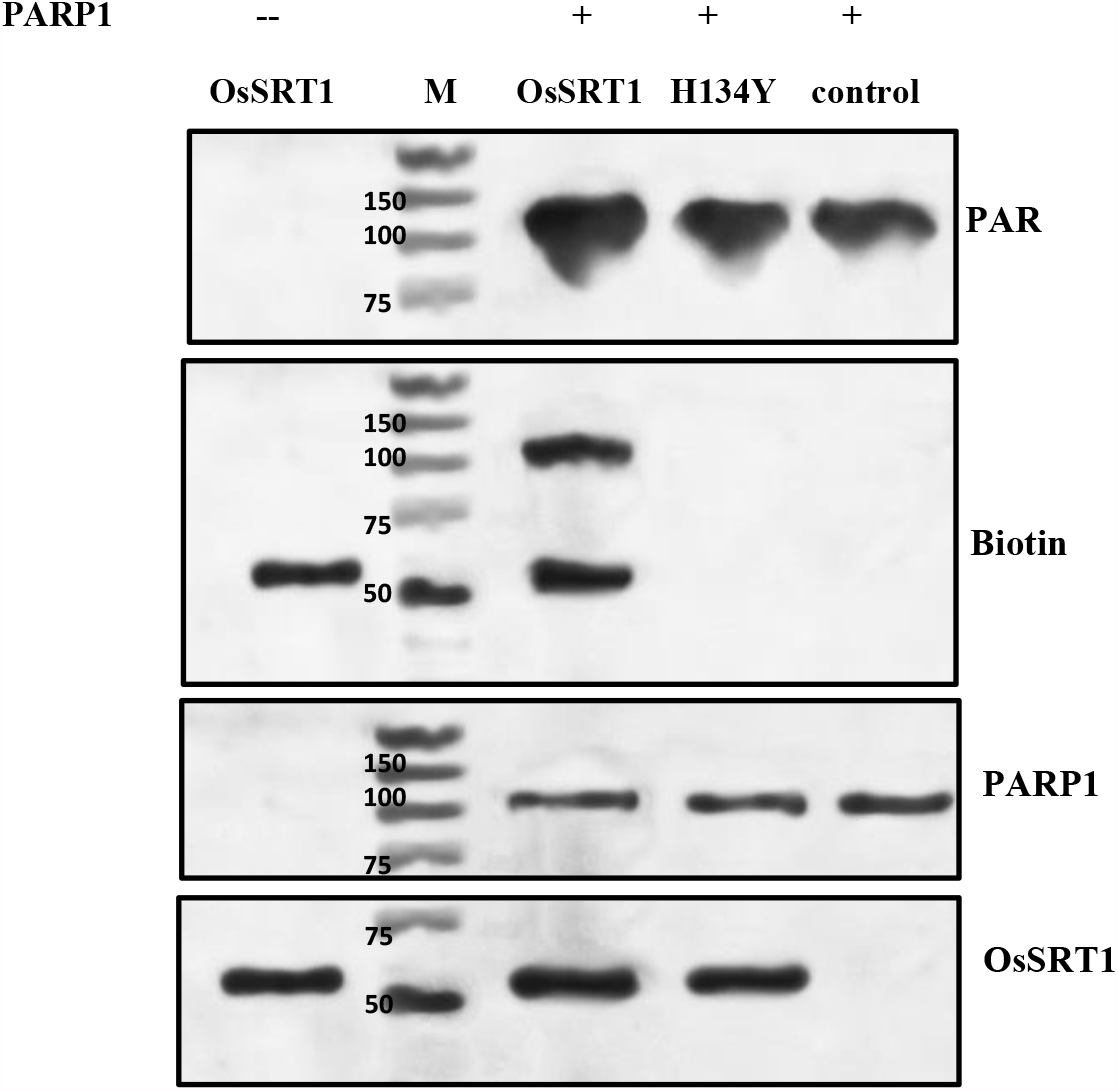
Effect of OsSRT1 ADP ribosylation on PARP1 activity: Western blot analysis using poly ADPr antibody showed that in presence of OsSRT1, the PARP1 activity increases. Addition of mutant H134Y or empty vector (control) did not modify the PARP1 activity as shown in lane 4 and 5.

**FigS5.**
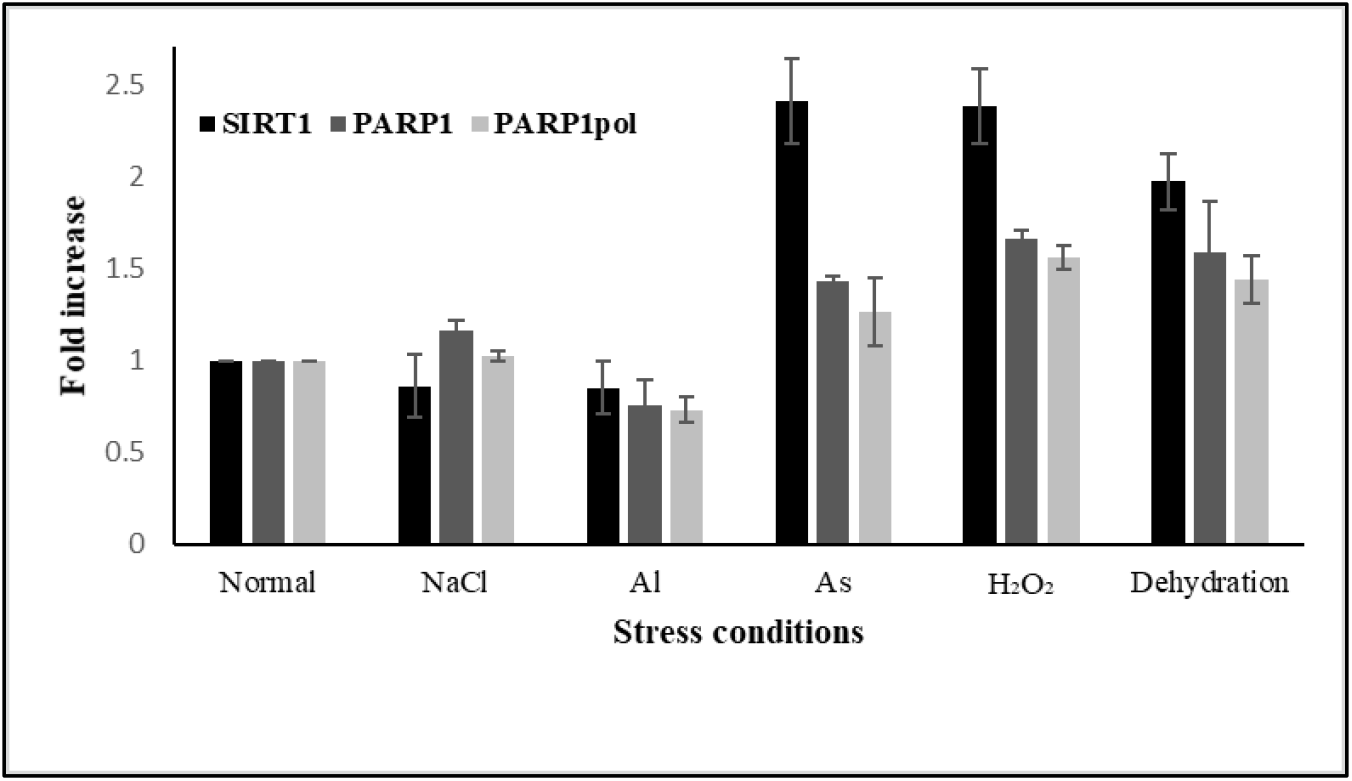
Relationship between the OsSRT1 expression and PARP1 activity under stress conditions: The nuclear protein extract was prepared from the leaves of the plants exposed to the different stress conditions as mentioned in the figure. Western blot analysis was performed using the specific antibodies (OsSRT1, PARP1 and PAR). Histone H4 in the nuclear extract was used as the internal control. All the data were normalized before plotting. The PARP1 enzyme activity increased with increase in expression of OsSRT1 under Arsenic, H2O2, and dehydration. It is possible that the OsSRT1 ADPr PARP1 to increase its activity.

